# Epigenetic Modulation of Host Immunity related genes in Pulmonary Tuberculosis: A Comprehensive Analysis of DNA Methylation Profiles in Peripheral Blood Mononuclear Cells

**DOI:** 10.1101/2025.01.16.633312

**Authors:** Ankit Kumar, Sumedha Sharma, Jyotdeep Kaur, Arnab Pal, Ashutosh Nath Aggarwal, Indu Verma

## Abstract

**Background:** In the last few years, there has been increasing evidence in the literature for promoter hypermethylation of various host genes following infection with various pathogens. Therefore, in the present study, the role of epigenetic reprogramming of host cells through DNA methylation was evaluated which indicates that the disease progression takes place through changes in host immune response related genes.

**Objective:** To elucidate the differential genome-wide DNA methylation profile of peripheral blood mononuclear cells in pulmonary tuberculosis (PTB) patients.

**Materials and methods:** To decipher the methylation profile of PBMCs in PTB patients whole genome bisulphite sequencing was performed using 4 DNA samples from each study group i.e. PTB, healthy controls and diseased controls. Briefly, the samples were subjected to bisulphite conversion followed by library preparation and whole genome bisulphite sequencing. The data then were analysed for differential methylation analysis followed by gene set enrichment analysis to select DMRs for validation by sanger sequencing. Further the GEOR2 online server was used to investigate the mRNA expression of selected genes associated with hypermethylated DMRs in promoter region.

**Results:** Differential methylation region analysis was performed for the following comparisons. TB v/s Diseased (Hypermethylated=1756; Hypomethylated=1886), TB v/s Healthy( Hypermethylated=2203; Hypomethylated=2466), Diseased v/s Healthy ( Hypermethylated=1656; Hypomethylated=3936). Further gene set enrichment analysis showed that the hypermetylated regions belonged to genes that are involved in immune responses mainly T cell functioning and T cell mediated immune processes. There were 8 DMR belonging to TAF8,FZD5, HLA-DRB, MIR483, PVRIG, SH2B2, ZAP70, TNFRSF13C simultaneously (n=10 from each study group). mRNA expression analysis with GEOR2 (Datasets used n= 12) revealed significant downregulation of TNFRSF13C, ZAP70, PVRIG Among the selected genes associated with hypermethylated DMRs in promoter region.

**(e) Conclusion:** The altered differential methylation profile of PBMCs from TB patients shows that on onset the active disease there occurs aberrant methylation of expression regulating regions that in turn results in decreased T cell functions and hence suboptimal immune responses in TB

## Introduction

*M. tuberculosis*, has survived for millennia in human communities, showcasing its adaptability and persistence. According to Hershkovitz et al., (2017) and Zaman (2010), there is evidence of tuberculosis (TB) in ancient human remains *(Hershkovitz, Donoghue et al. 2008), (Zaman 2010)*, shedding light on intriguing understanding into the long-standing connection that exists between this pathogen and the host *(Adigun and Singh 2023)*. M.tb is very well known for its ability to avoid detection by the host immune system and stay out of sight for a long time and flourishing within the host as latent tuberculosis. Nonetheless, the M.tb has evolved a defense mechanism of its own, concentrating on evading our immune system and thus giving rise to the clinical manifestations *(Chandra, Grigsby et al. 2022).*The immune system is known to play an important role in providing defense against various infections. In the case of TB, suppression of the immune system which favors pathogen survival and efficient proliferation inside the host is well known (Brett, Dulong et al. 2020). During the entire life cycle of Mtb from inhalation of a few bacteria in the lung to the generation of large number of bacilli in the airway by liquefaction of granuloma, there is a constant battle between Mtb and immune system (de Martino, Lodi et al. 2019). Although, immune system is able to control infection in a majority of infected individuals, a small proportion of them loses this battle and develops active TB. Currently, the mechanism of immune defense against mycobacteria is not clearly known, however, our understanding has considerably increased due to several studies performed in the past decade. These studies have demonstrated that both innate and adaptive immune systems are involved in providing immunity against Mtb. The outcome of the immune response against Mtb depends upon multiple factors, including crosstalk among innate and adaptive immune pathways (Zhang, Gao et al. 2023). Epigenetic changes such as DNA methylation are reported to be modulated in infectious diseases. It is not very clear at this moment that how DNA methylation gets altered at the time of infection and during progression of the disease. It is speculated to happen either due to inflammatory reactions or by direct involvement of bacterial factors. *Helicobacter pylori* is one such pathogen that is known to modulate DNA methylation patterns by inducing inflammation. There are other pathogens like *Porphyromonas gingivalis*, *Escherichia coli* and *Chlamydophila pneumonia* which cause the alteration in methylation of genes specifically involved in the immune responses (Murata, Azuma et al. 2007, Huang and Berger 2008, Tolg, Sabha et al. 2011, Yin and Chung 2011, Harouz, Rachez et al. 2014) . Alterations in DNA methylation profile of host immune cells is known in TB. This altered methylation is suggestive of diversion of the immune response to favor the intracellular survival and proliferation of M.tb inside the host.

## Results

### Host genome is hypermethylated at global level in active TB patients

To check if there is any difference in overall levels of methylation (Global methylation) in TB patients when compared to diseased controls and healthy controls, COBRA (Combined Bisulfite Restriction analysis) assay was performed. *LINE-1* is a repetitive element and is known to be hotspot for methylation throughout the human genome hence percent methylation present in *LINE-1* is a direct indicator of overall methylation status throughout the genome. Because the original pool of DNA utilized for bisulfite conversion comprises both methylated and non-methylated cytosines at the genomic locus, bisulfite conversion produces a mixed population of DNA fragments of different sizes as explained in the illustration (fig. 1a).

**Fig. 1.**
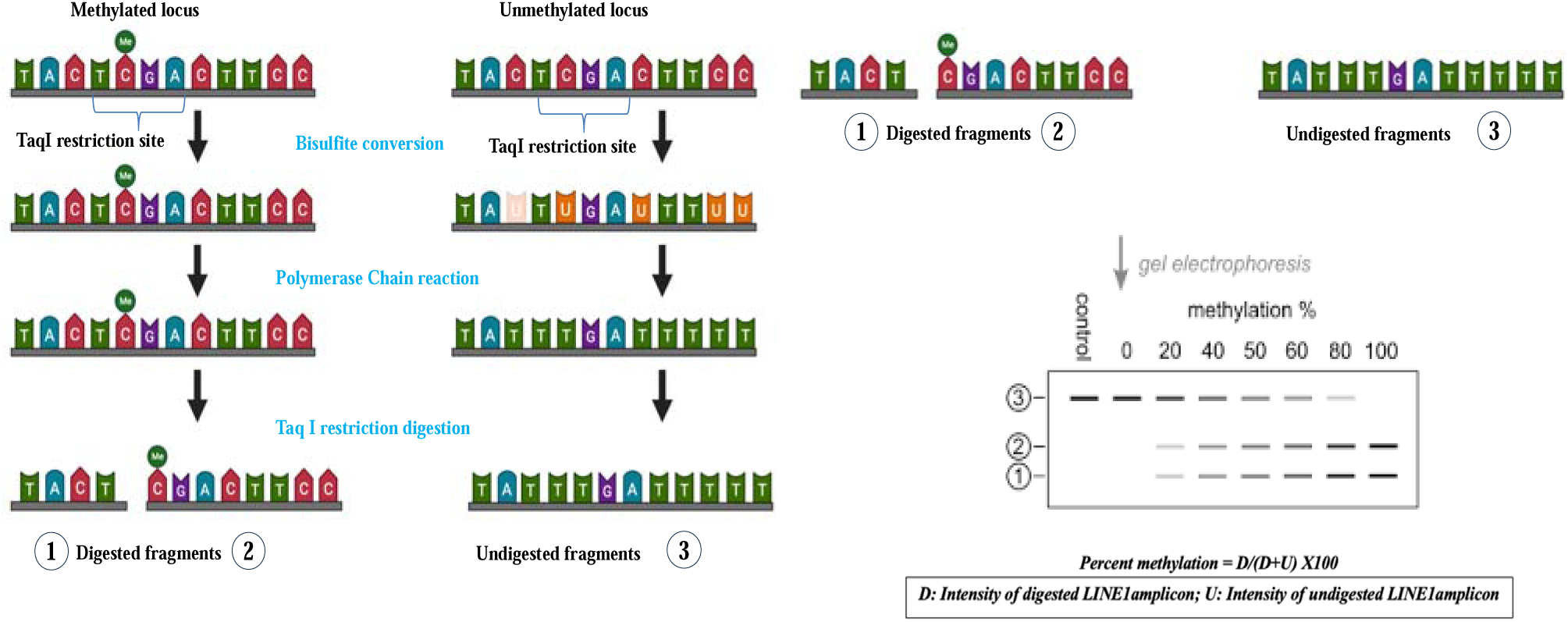
a: A schematic illustration of COBRA assay (generated using Bio render)

**Fig. 1.**
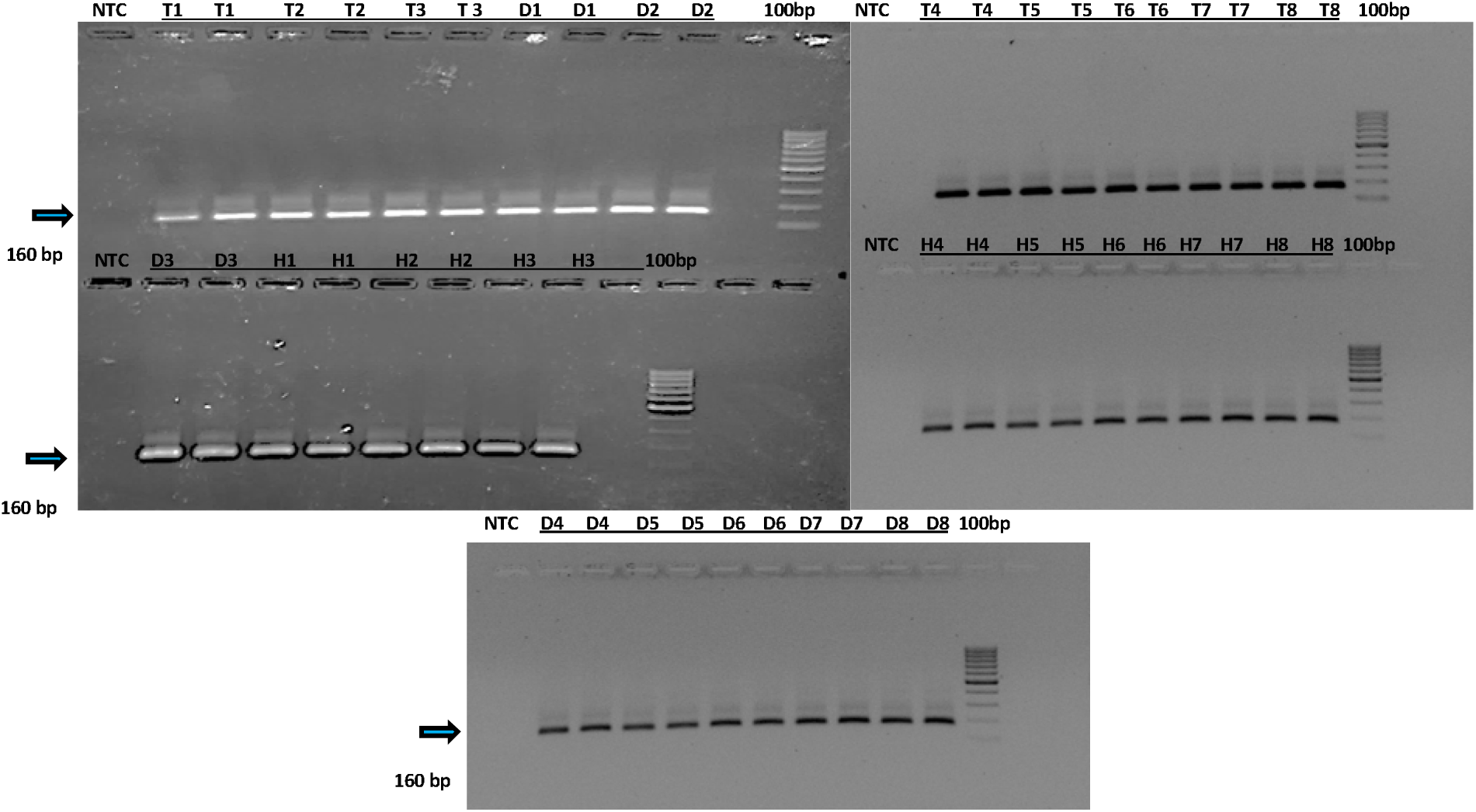
b Gel pictures of amplified LINE-1 gene (160 bp) on 2% agarose gel

**Fig. 1.**
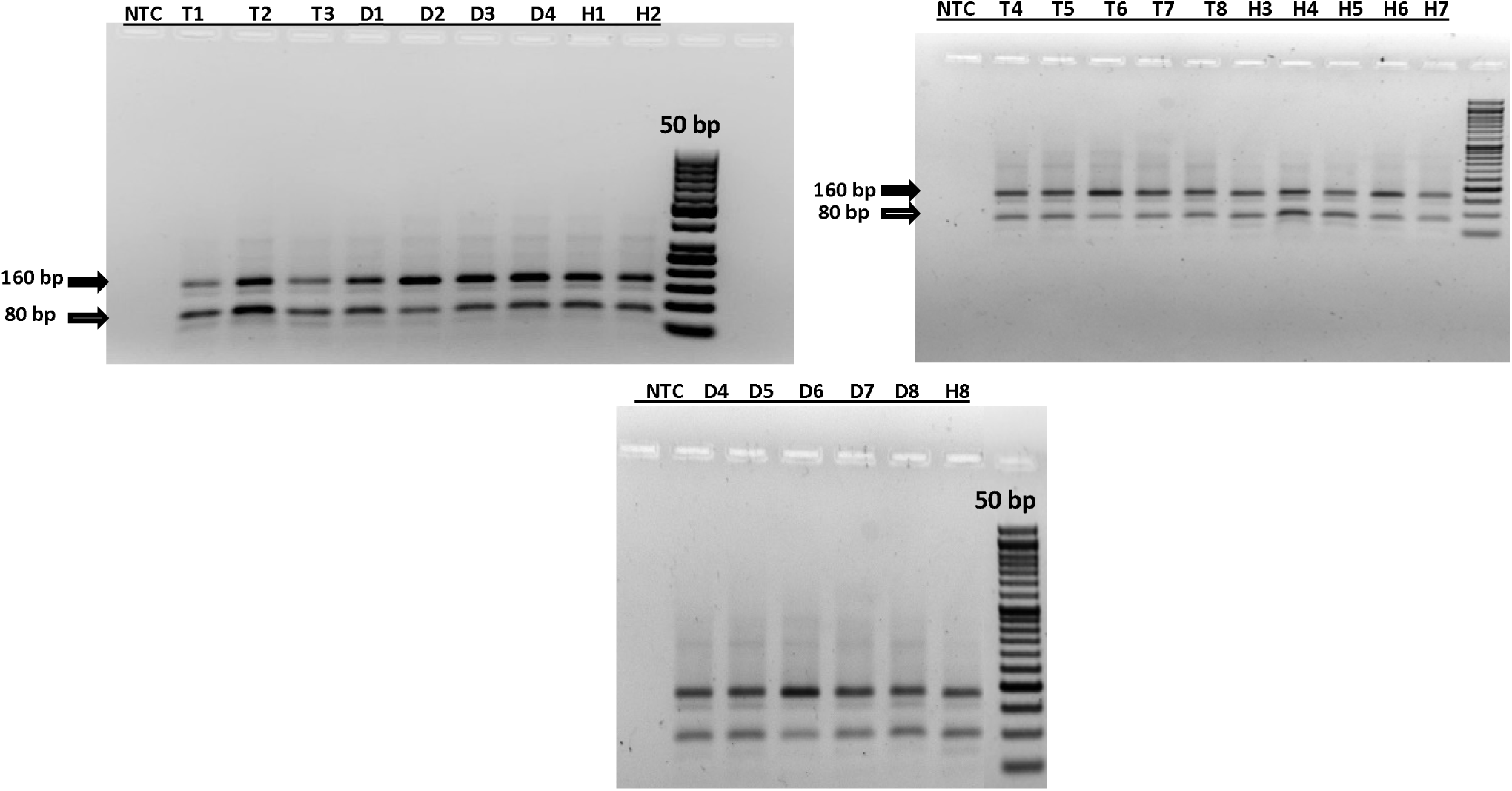
c: Gel pictures of amplified LINE-1 gene after digestion with TaqI restriction digestion on 2% agarose gel

**Fig. 1.**
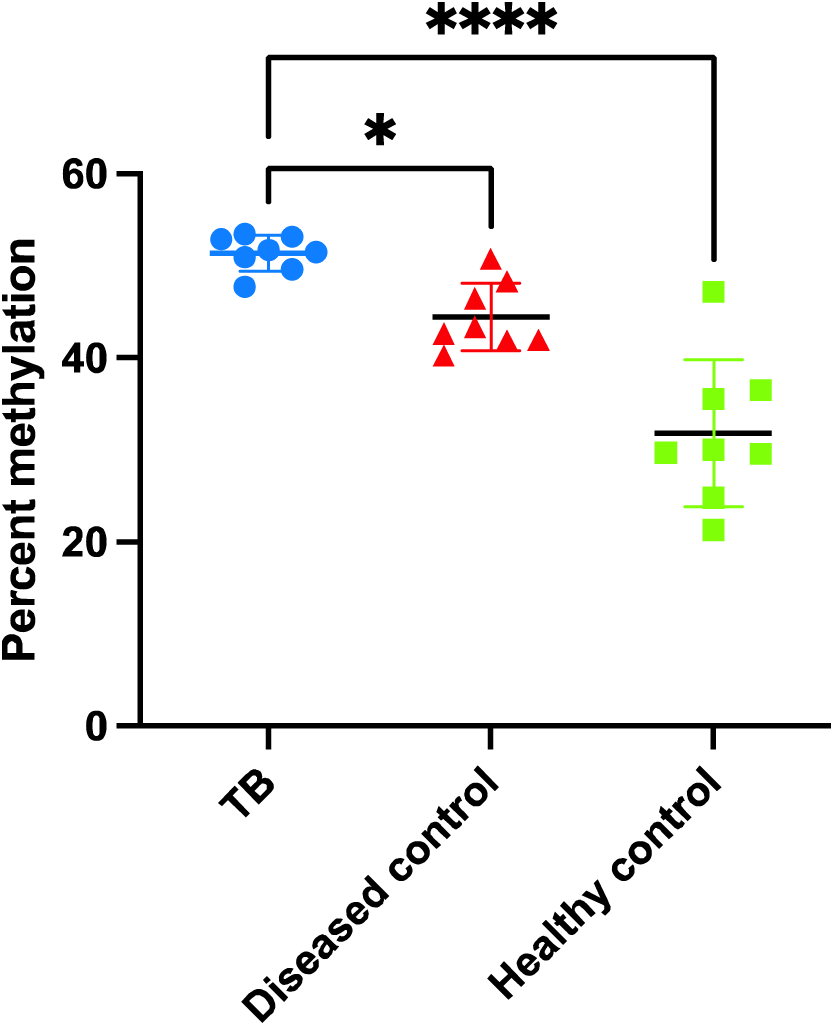
d: Dot plot graph of percent methylation of three study groups,values are represented as Mean ± SD; *p≤ 0.05, **p ≤0.01, ***p≤ 0.001, ****p≤ 0.0001 , using Kruskal Wallis test followed by Dunn’s multiple comparison test

For this assay 8 samples from each study group i.e. TB, healthy and diseased controls were used. Following the three sequential stages of the assay, the PCR amplicon for each sample was run on 2% agarose gel to verify the amplification of *LINE-1* genomic sequence and after digestion with TaqI (fig. 1b). Densitometry analysis showed a significant difference in levels of percent global methylation among the three study groups being highest in TB group (51.38 ±1.950) when compared to healthy (31.83 ±7.967) and diseased controls (44.47±3.658) fig. 1c. This further prompted an elaborated exploration of methylation changes of host at genome wide level.

### TB patients have a differentially altered methylome

To decipher the differential genome-wide DNA methylation profile of peripheral blood mononuclear cells in pulmonary tuberculosis (PTB) patients, the genome wide methylation profile was analyzed within PBMCs obtained from anti-tuberculous treatment naive TB patients in comparison to diseased and healthy controls (n=4 each study group) using whole genome bisulfite sequencing (WGBS). The sequencing libraries were prepared using 200ng of genomic DNA followed by bisulfite treatment, addition of indexing adapters and the purification. The purified libraries were quantified using qPCR and then sequenced on Novaseq™ (Illumina) at 30X coverage, 2x150bp reads/sample. Up to 75% of the sequenced bases were of Q30 value >85. Sequenced data was processed to generate FASTQ files and proceeded with differential methylation region analysis for the following comparisons: TB v/s diseased and TB v/s healthy. Differentially methylated regions (DMRs) were filtered with percent methylation cut off of ±30%. Analysis revealed 3642 DMRs for TB vs diseased controls comparison of which 1756 were hypermethylated and 1886 hypomethylated DMRs whereas for TB vs healthy controls comparison there were 4667 DMRs, of which 2464 were hypermethylated and 2203 were hypomethylated. Data was also analyzed to check the number of DMRs that falls into promoter region (Differentially methylated promoters), and it was found that for TB vs diseased controls comparison there were 219 DMRs in promoter regions of which 77 were hypomethylated and 142 were hypermethylated whereas for TB vs healthy controls comparison there were 359 DMRs in promoter regions of which 254 were hypomethylated and 105 were hypermethylated (Fig. 2a & 2b).

**Fig. 2.**
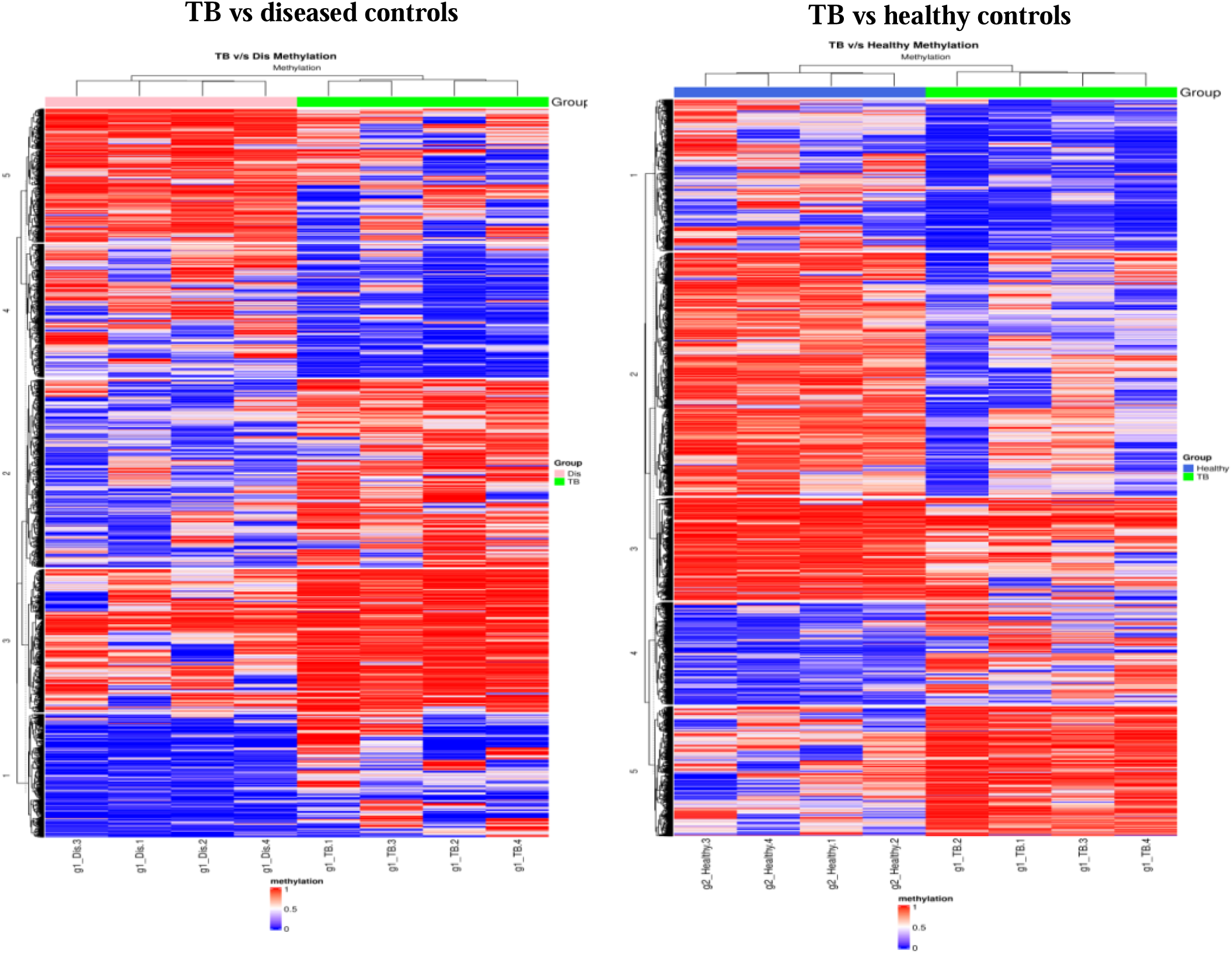
a: Heat Maps of differential methylation profiles of PBMCs from TB, Diseased and healthy controls

**Fig. 2.**
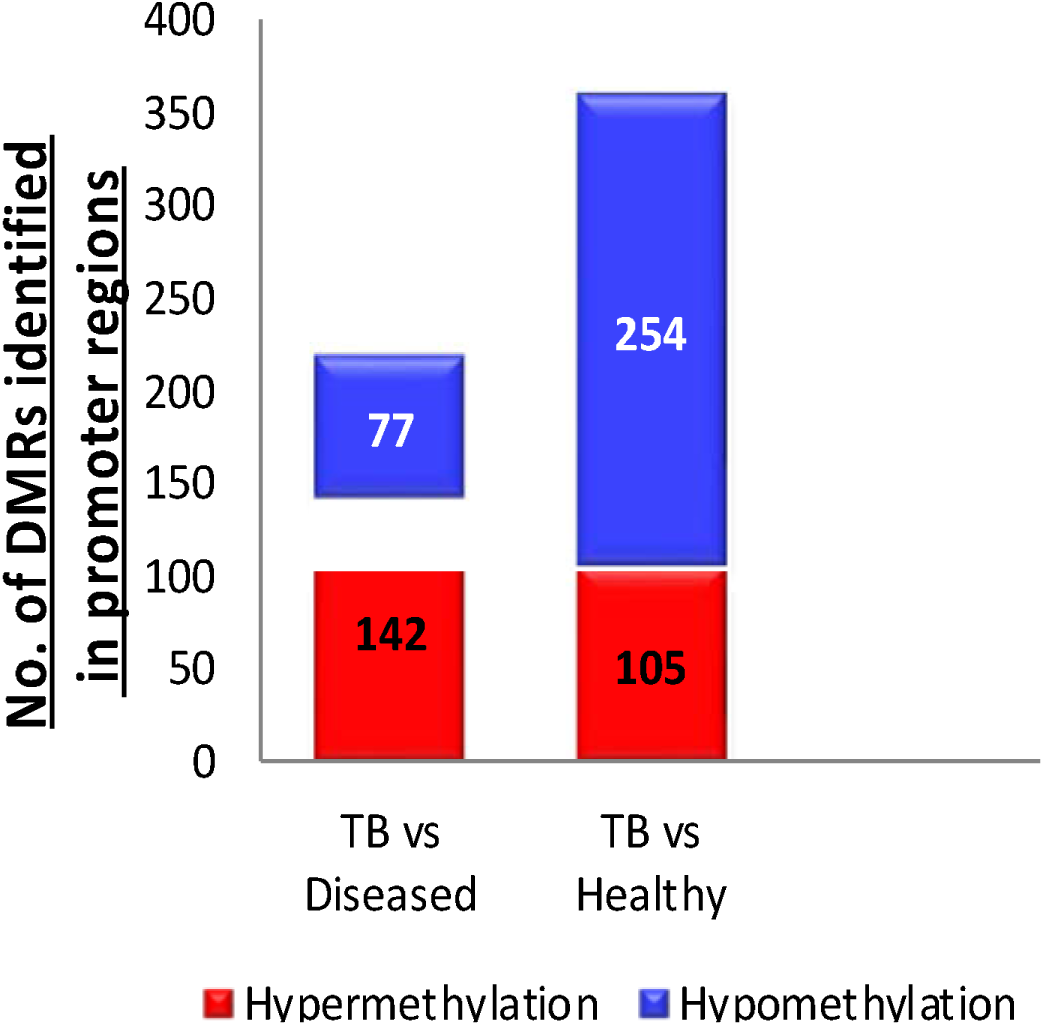
b: Graph depicting DMRs identified in the promoter region for both comparisons

**Fig. 2.**
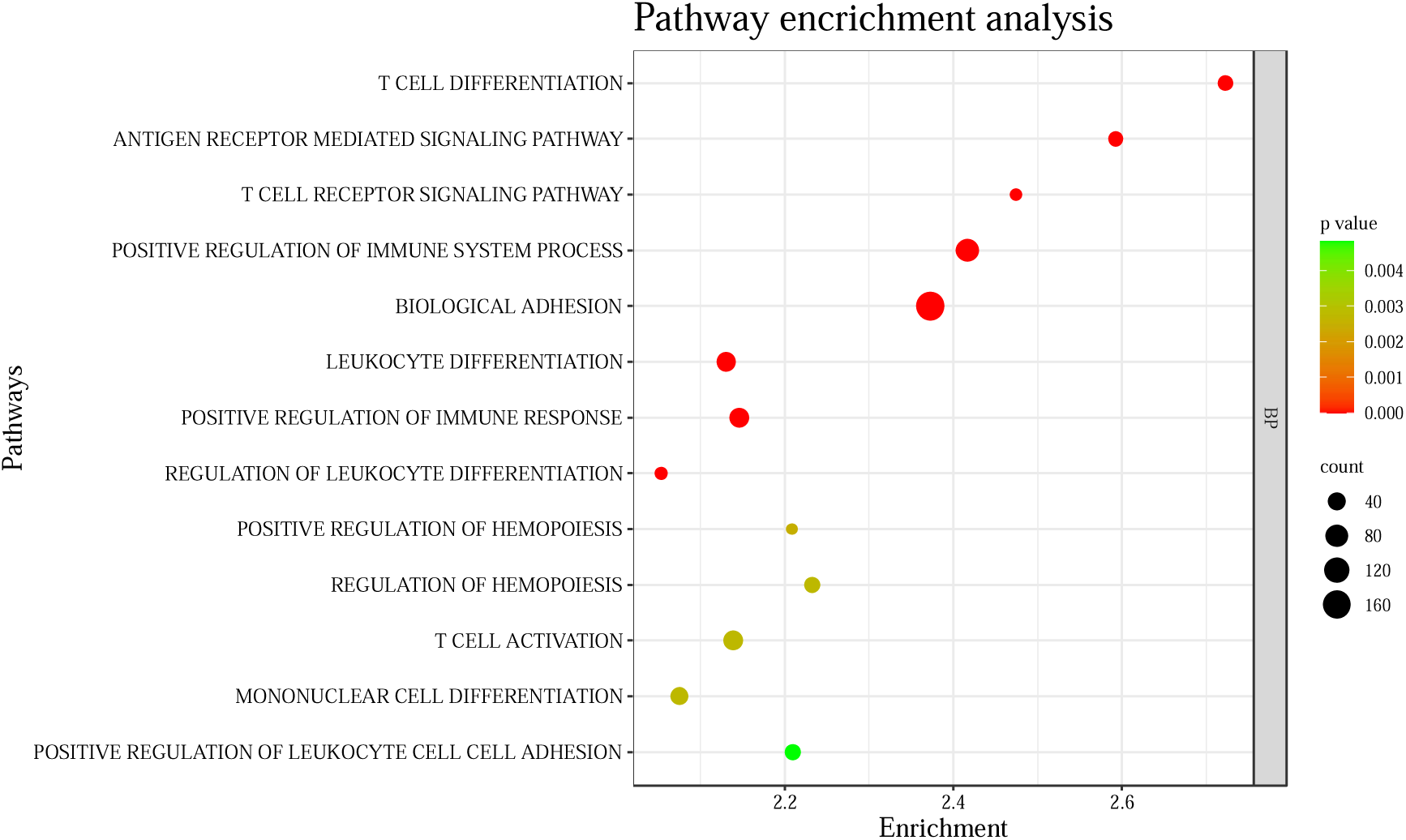
c: Bubble plot of significantly positively/ negatively enriched terms (generated using SR plot online server) for TB vs Healthy controls comparison; count depicts the number of genes in a dataset and p value is indicative of statistical significance of enriched gene sets.

**Fig. 2.**
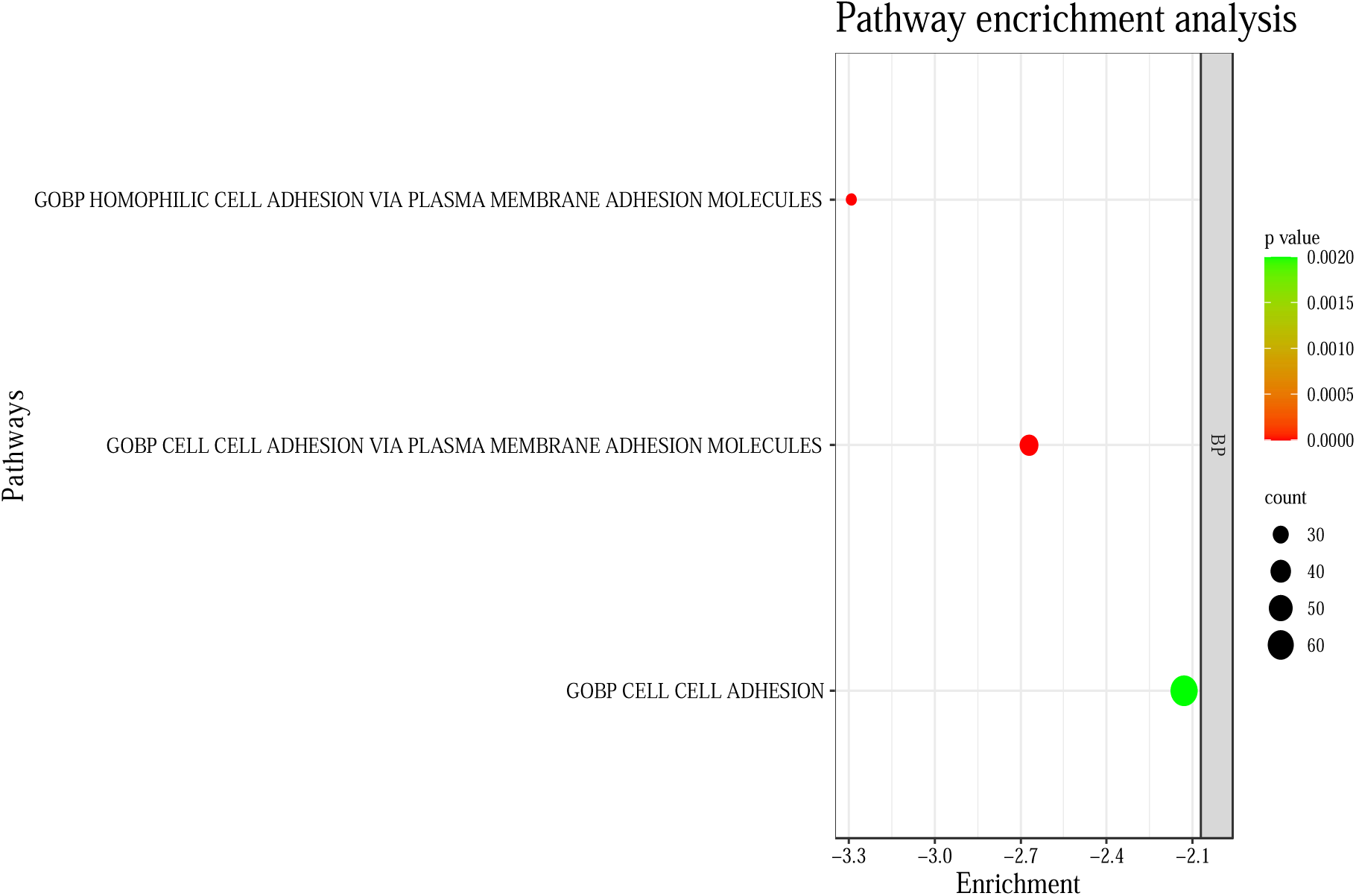
d: Bubble plot of significantly positively/ negatively enriched terms (generated using SR plot online server) for TB vs Diseased controls comparison; count depicts the number of genes in a dataset and p value is indicative of statistical significance of enriched gene sets.

### T cell immune responses related genes are specifically hypermethylated in TB patients

Gene set enrichment analysis (GSEA) revealed positive enrichment of 13 gene sets corresponding to pathways involved in host immune responses and were as follows: T cell differentiation, antigen receptor signaling pathway, T cell receptor signaling pathway, positive regulation of immune system process, leukocyte differentiation, positive regulation of immune system process, positive regulation of immune system, biological adhesion, leukocyte differentiation, positive regulation of immune response, regulation of leukocyte differentiation, positive regulation of hemopoiesis, regulation of hemopoiesis, T cell activation, mononuclear cell differentiation and positive regulation of leukocyte cell cell adhesion in TB vs healthy controls comparison (Fig. 2c), however there were no significantly negatively enriched pathways as revealed by the analysis. In TB vs diseased controls comparison there was no significantly positively enriched pathway, however significantly negatively enriched pathways included homophilic adhesion via plasma membrane adhesion molecules, cell cell adhesion via plasma membrane adhesion molecules and cell cell adhesion (fig. 2d).

There was a total of 30 hypermethylated DMR associated genes that were involved in top three significant positively enriched pathways associated with immune responses. These 30 hypermethylated DMR associated genes were further analyzed for methylation status of their promoter region. This analysis revealed 7 such genes that were having hypermethylated DMRs in their promoter regions and were common to immune response pathways (T cell related immune responses) (table 1) .

**Table 1:** List of DMRs selected for Validation.

| **Sr. No.** | **Location of DMR** | **CpGs** | **Differential methylation** | **Length bp** | **Promoter name** | **Function** | **Pathway involved** |
| --- | --- | --- | --- | --- | --- | --- | --- |
| **1** | Chr6:42050104 to 42050153 | 5 | 0.42 | 149 | TAF8 | TATA box binding protein associated factor 8 | T cell activation , T cell differentiation, T cell receptor signalling |
| **2** | Chr6:32590191 to 32590598 | 5 | 0.43 | 407 | HLA-DRB1 | Major Histocompatibility Complex, Class II, DR Beta 1 | T cell activation |
| **3** | Chr11:2134969 to 2135129 | 5 | 0.38 | 160 | MIR-483 | MicroRNA 483 | T cell receptor signalling pathway |
| **4** | Chr7:100217675 to 100217864 | 5 | 0.31 | 189 | PVRIG | Poliovirus receptor related immunoglobulin domain containing | T cell activation , T cell differentiation |
| **5** | Chr7:102284140 to 102284259 | 5 | 0.35 | 119 | SH2B2 | SH2B Adaptor Protein 2 | T cell activation , T cell differentiation, T cell receptor signalling |
| **6** | Chr2:97712907 to 97713347 | 5 | 0.34 | 440 | ZAP70 | Zeta-chain-associated protein kinase 70 | T cell activation |
| **7** | Chr22:41927108 to 41927208 | 5 | 0.30 | 100 | TNFRSF13C | TNF Receptor Superfamily Member 13C | T cell activation , T cell differentiation, T cell receptor signalling |

### Hypermethylation of DMRs in non CpG island context in T cell immune response related genes

Based upon findings of WGBS and GSE analysis, the 7 identified genes were selected to further determine the specific methylation location in promoter region. For this the location coordinates of hypermethylated DMRs corresponding to the 7 selected genes were loaded as input in UCSC genome browser one by one and location of methylated CpGs were identified. It was revealed by this analysis that there is a non-canonical CpG methylation in the promoter region of all the selected 7 genes and four genes ( *TAF8, HLA-DRB1, PVRIG, ZAP-70*) had this non-canonical CpG methylation in the proximal enhancer region of the promoter sequence as shown in the fig. 3a.

**Fig. 3.**
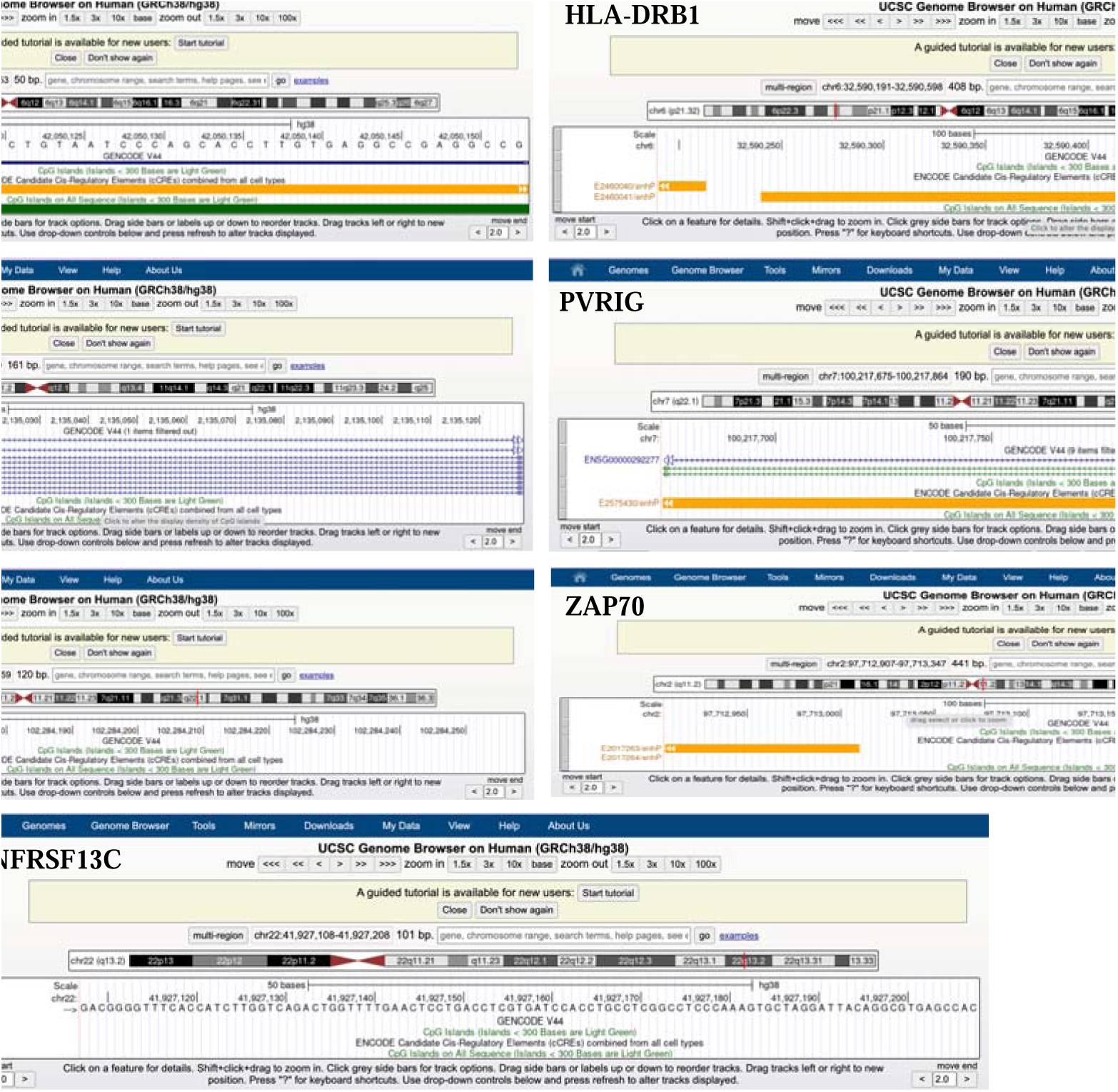
a: Pictures representing results from UCSC genome browser analysis for each selected DMRs

**Fig. 3.**
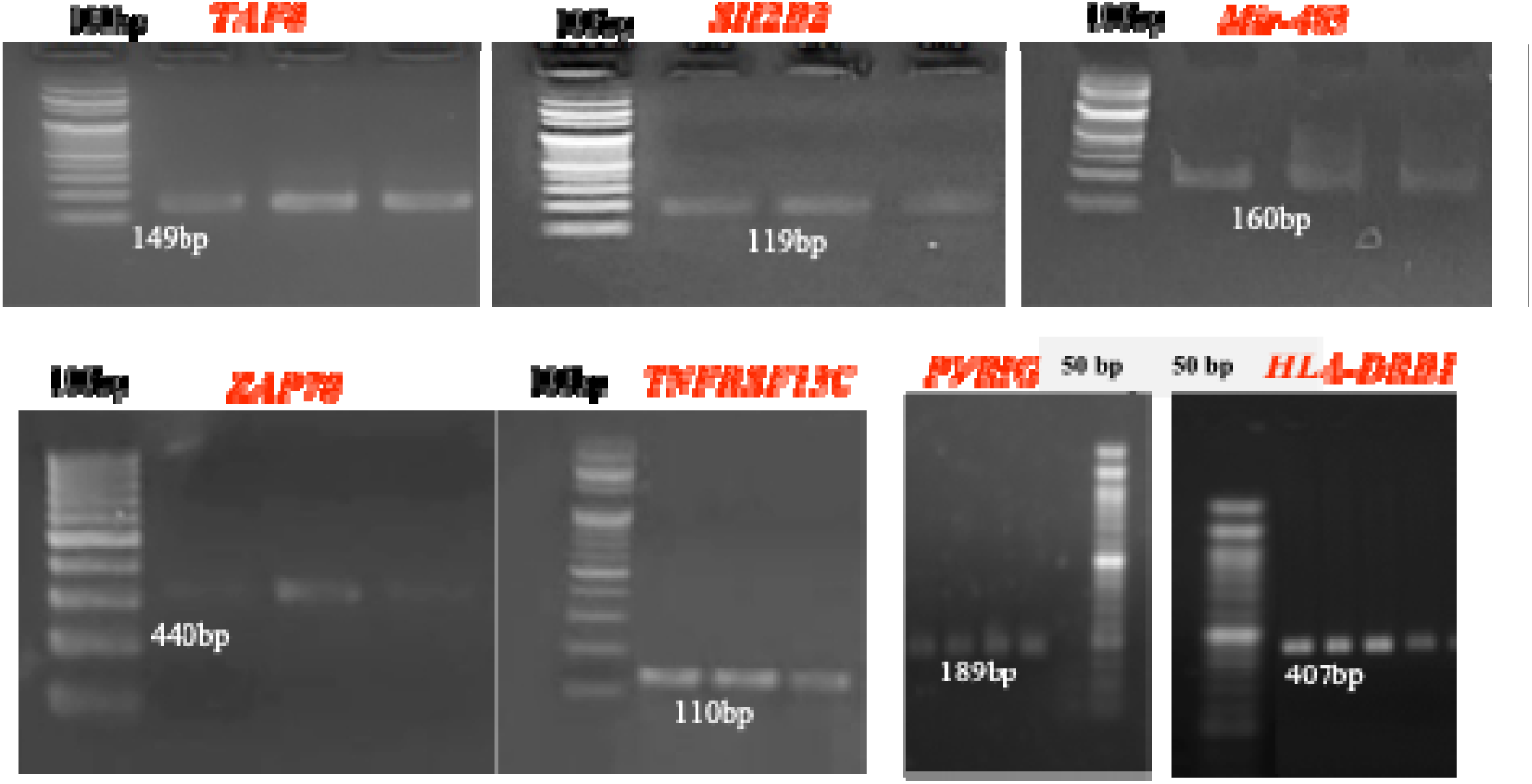
b: Representative gel image of selected amplified DMRs using methylation specific primers.

**Fig 3.**
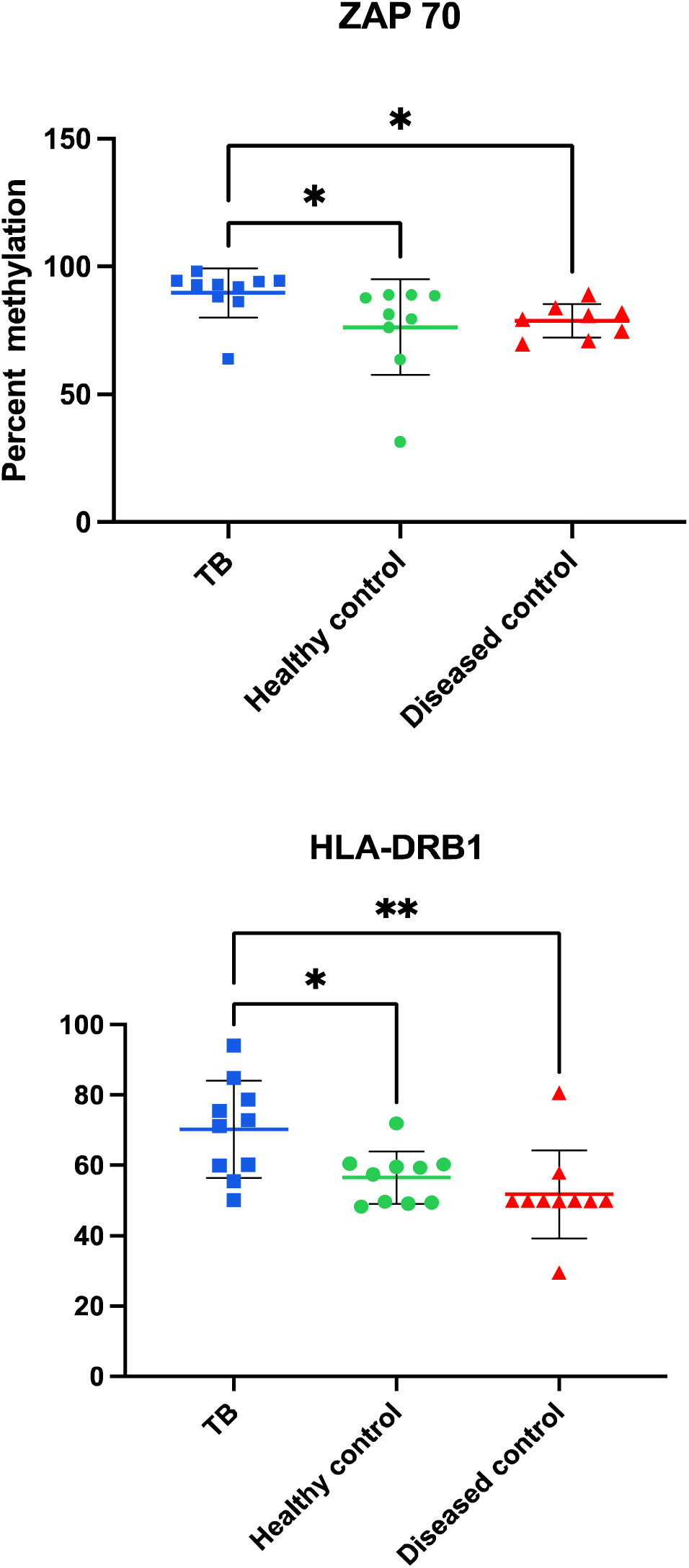
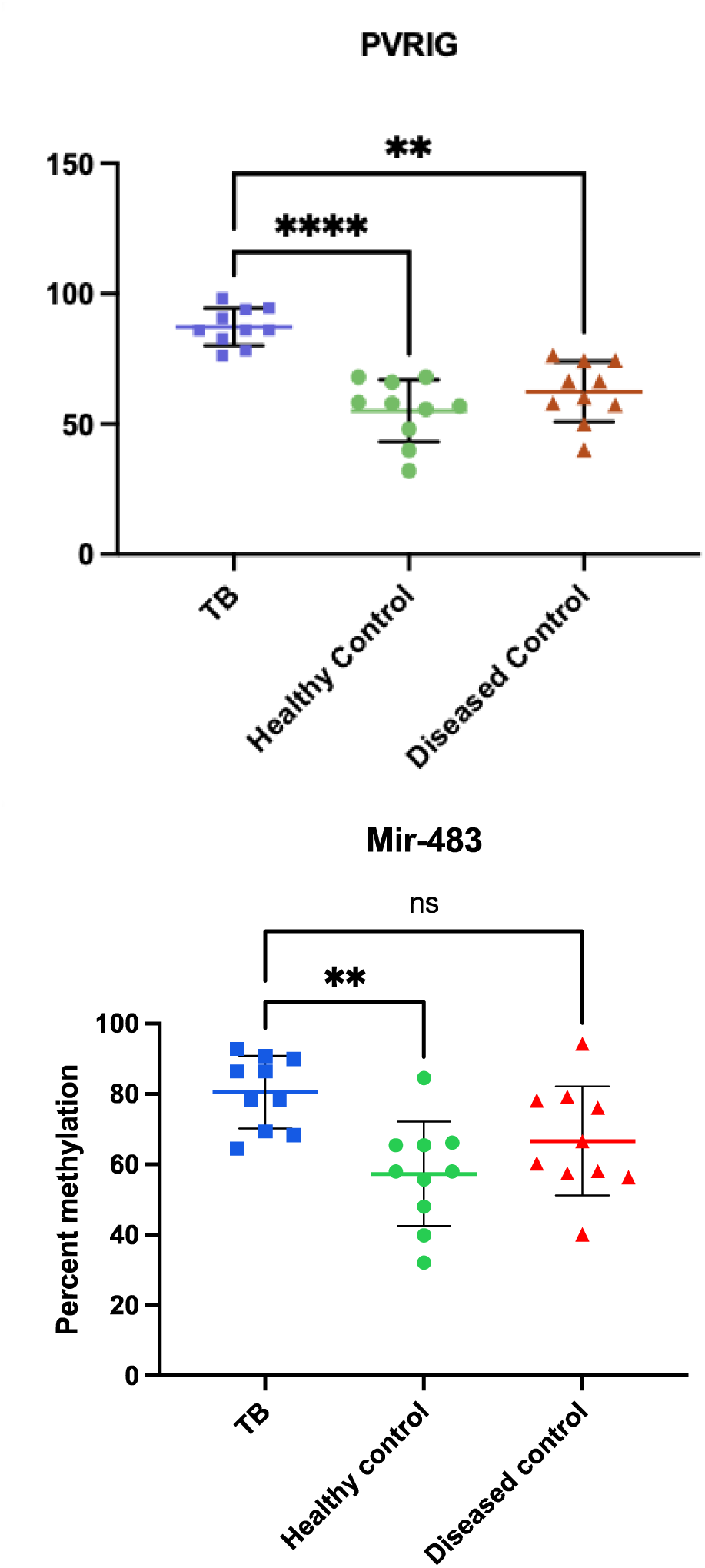
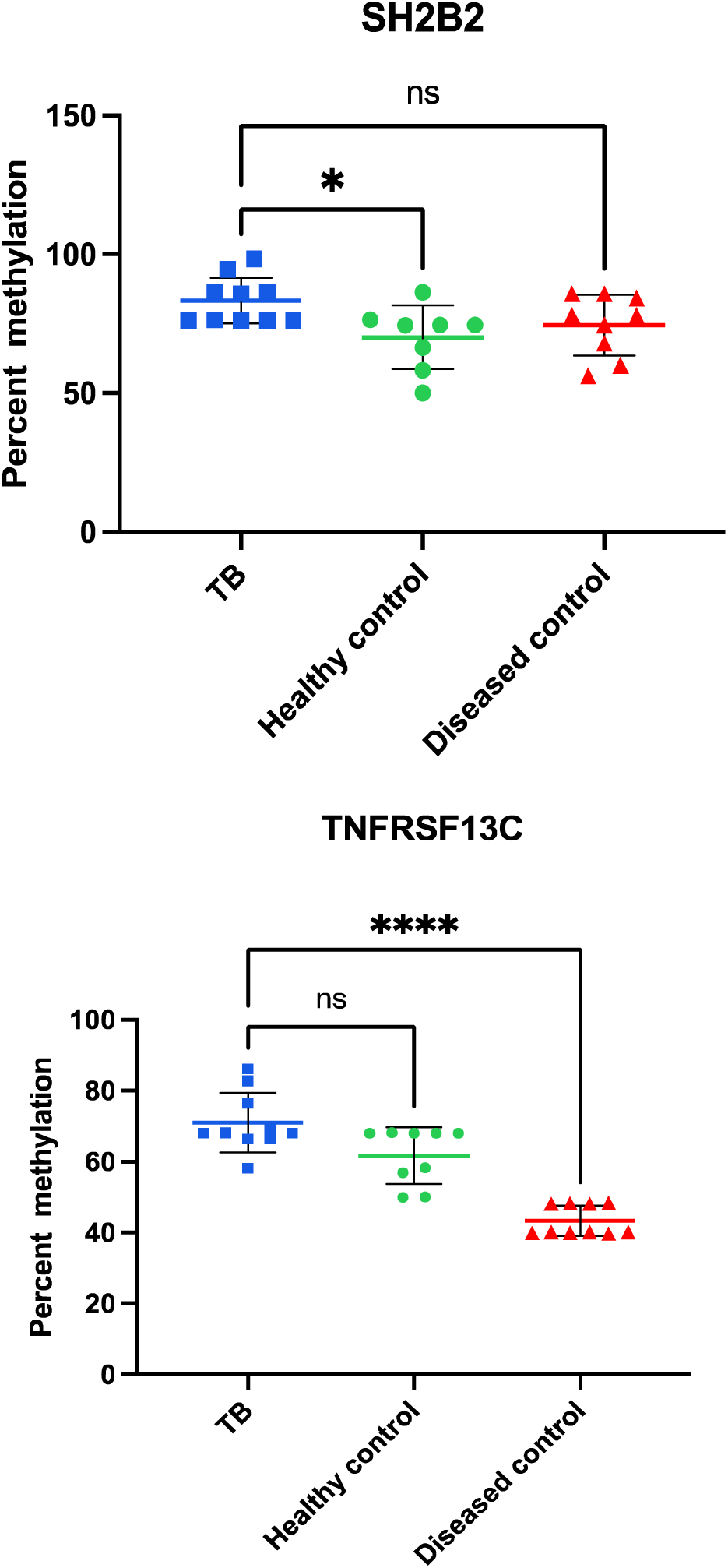
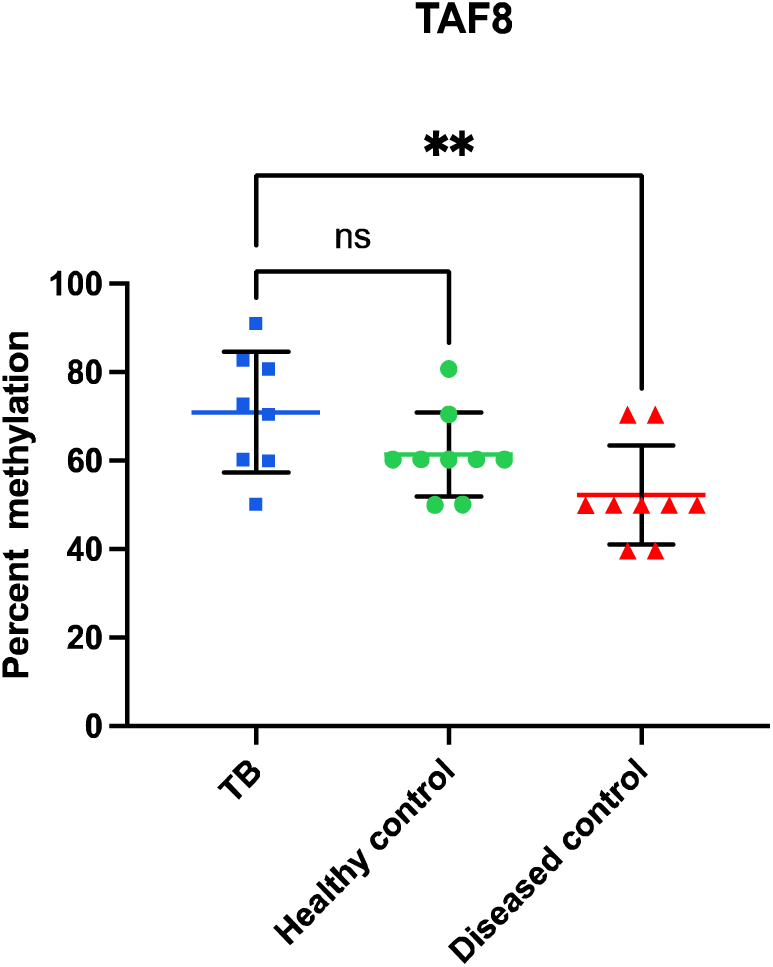
c: Validation of differentially methylated regions identified by WGBS from PBMCs of TB patients in comparison to diseased controls and healthy controls using indirect bisulfite sequencing: Dot plot of individual % methylation values of each sample for each DMR, data represented as Mean ± SD; *p≤ 0.05, **p ≤0.01, ***p≤ 0.001, ****p≤ 0.0001 , using Kruskal Wallis test followed by Dunn’s multiple comparisons test ;n=10 each for TB, healthy and diseased control group.

**Fig 3.**
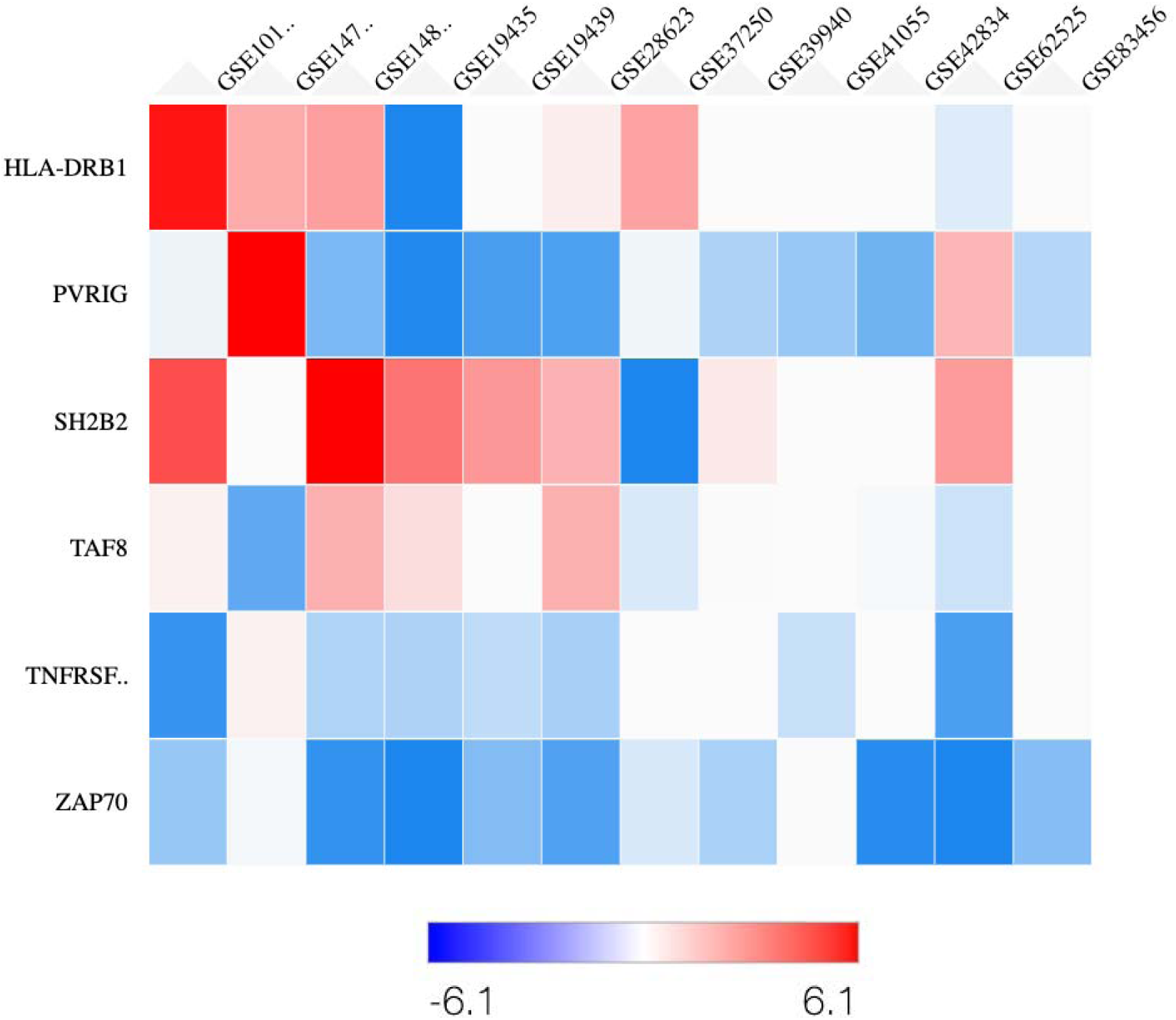
d: Heat map depicting gene expression (log2 Fold change) of seven selected genes in twelve datasets retrieved from GEO database in silico gene expression analysis.

Based upon the WGBS, GSEA and UCSC genome browser analysis, 7 genes (*TAF8* (149bp), *SH2B2* (173bp), *Mir-483* (180bp), *ZAP70* (329bp), *TNFRSF13C* (125bp), *PVRIG* (181bp), and *HLA-DRB1* (300bp) were selected for validation (Fig. 3b). For validation of WGBS data indirect bisulfite sequencing was used and 10 samples from each study group were used. PCR amplified amplicons (supplementary table 1, 2 and 3)of the above mentioned DMRs were taken further for sanger sequencing by dye-terminator sequencing. Sequenced samples were analyzed, and peak height was calculated for Cytosine(C) and Thymine(T) in the DNA sequence for calculation of C:T ratio (supplementary figure 1).

Out of the seven selected genes associated with DMRs, the methylation status of DMRs of *TAF8* (TB—70.98 ±13.66, disease controls—52.21 ±11.17) and *TNFRSF13C* showed significant hypermethylation in TB patients when compared to diseased controls however the difference was not significant in comparison to healthy controls (TB—76.77 ±9.28, disease controls—41.78 ±6.04). DMRs associated with *SH2B2* and *Mir-483* genes were found to be significantly hypermethylated in comparison to healthy controls whereas the difference was not significant in comparison to diseased controls. In case of *HLA-DRB1* (TB—70.30 ±13.83) was significantly higher as compared to both healthy (51.76 ±12.43) and disease controls (56.56 ±7.48). In *ZAP70* and *PVRIG* genes also, significantly hypermethylation was observed in TB patients in comparison to both the study groups, healthy as well as diseased controls (Fig. 3c).

To further validate the findings of WGBS, *in-silico* screening of gene expression of the seven selected genes was done using GEO2R, *In-sillico* gene expression analysis of selected genes revealed that gene expression of PVRIG, ZAP70 and TNFRSF13C genes are downregulated in majority of the datasets (Fig. 3d). This further strengthens the finding of WGBS because the DMRs associated with these genes have methylation in promoter proximal cis regulatory elements and thus can downregulate gene expression to a greater extent.

## Discussion

It is well known that M.tb has the ability to adapt itself in various host microenvironments by altering its gene/protein expression profiles and these changes in transcriptional or proteome profiles may further be associated with disease pathogenesis (Sharma, Ryndak et al. 2017, von Both, Berk et al. 2018). Although there are few studies where mycobacterial proteins have been shown to affect the DNA methylation of host immune pathways genes , however, there is no report on the role of *in vivo* expressed mycobacterial proteins during active TB in patients in the modulation of host immune response particularly through epigenetic changes (Sharma, Upadhyay et al. 2015, Yaseen, Kaur et al. 2015). The findings presented in this study shed light on the intricate interplay between *Mycobacterium tuberculosis* (Mtb) infection and the epigenetic landscape of host immune cells, particularly focusing on DNA methylation patterns. Tuberculosis (TB) remains a significant global health concern, and understanding the mechanisms by which Mtb modulates host immunity is crucial for developing effective therapeutic interventions. The findings of this study provide significant insights into the epigenetic modulation of host immune responses in pulmonary tuberculosis (TB). The observed global hypermethylation in active TB patients suggests a systemic alteration in the epigenetic landscape of the host, potentially contributing to immune evasion by M.tb.

LINE-1 elements, used as markers for global methylation, showed significantly higher methylation levels in TB patients (51.38%) compared to healthy (31.83%) and diseased controls (44.47%). This global hypermethylation could be indicative of an overall repressive chromatin state, which might hinder the expression of genes critical for effective immune responses. Previous studies have highlighted similar trends in other infectious diseases. For instance, *Helicobacter pylori* and *Porphyromonas gingivalis* are known to induce DNA methylation changes that affect immune response genes, facilitating chronic infection and immune evasion (Murata, Azuma et al. 2007, Huang and Berger 2008, Tolg, Sabha et al. 2011, Yin and Chung 2011, Harouz, Rachez et al. 2014). The hypermethylation observed in TB patients aligns with these findings, suggesting that M.tb may also exploit host epigenetic mechanisms to modulate immune responses. In previous reports on DNA methylation investigation in TB have been investigated in comparison to healthy controls, however only limited studies used diseased controls (respiratory infections other than TB) in this context. So, our study provides true and specific DNA methylation landscape, which opens new gates for biomarker discovery as well as host based therapeutic interventions for TB.

The whole-genome bisulfite sequencing (WGBS) data revealed significant differentially methylated regions (DMRs) in TB patients, particularly in genes involved in immune responses. Notably, genes such as *PVRIG*, Z*AP70*, and *TNFRSF13C* showed hypermethylation in their promoter regions. These genes play critical roles in T cell signaling and immune activation. Hypermethylation in these regions likely leads to their transcriptional repression, as supported by in-silico gene expression analyses showing downregulation of these genes in TB patients. The importance of T cell-mediated responses in controlling TB infection is well-documented. Studies have shown that effective T cell responses are crucial for containing M.tb infection and preventing its dissemination (Cooper 2009, Chang, Wherry et al. 2014) . The observed hypermethylation and subsequent downregulation of key T cell response genes in TB patients suggest a potential mechanism by which M.tb evades immune surveillance, thereby facilitating chronic infection and disease progression.

Thirteen gene sets related to immune responses were positively enriched in TB patients compared to healthy controls. These included pathways crucial for T cell differentiation and activation, leukocyte differentiation, and immune system regulation. The absence of significantly negatively enriched pathways in TB patients highlights a selective epigenetic repression of immune-activating pathways, possibly contributing to an impaired immune response. Epigenetic regulation of immune genes through DNA methylation has been observed in various contexts. For example, in chronic infections, epigenetic modifications can lead to a sustained suppression of immune responses, aiding pathogen persistence (Cuddapah, Barski et al. 2010, DiSpirito and Shen 2010)

The validation of methylation status through indirect bisulfite sequencing and in-silico gene expression analysis confirmed the hypermethylation of key immune genes. Notably, genes like HLA-DRB1, ZAP70, and PVRIG showed significant hypermethylation in TB patients, which correlated with their decreased expression levels. These results provide robust evidence supporting the role of DNA methylation in modulating immune gene expression during TB infection.

## Materials and methods

### Study subjects and sample collection

For studying the global DNA methylation levels and genome-wide DNA methylation pattern of peripheral blood mononuclear cells (PBMCs) following groups of HIV negative patients were recruited from the DOTS center, New OPD of PGIMER, Chandigarh after approval from institutional ethics committee vide no. INT/IEC/2019/001596 and informed consent was taken from each subject prior to sample collection.

a) Patients with pulmonary tuberculosis (n=10):

PTB patients having the clinical and radiological (chest X-ray) evidence of TB along with high AFB positivity(++/+++) in direct sputum smear microscopy or GeneXpert positive sputum were included in this group.

b) Diseased Controls (n=10):

Patients having other infections of the upper respiratory tract, lower respiratory tract and lung cancer with sputum expectoration were included in this group.

c) Healthy controls (n=10):

Healthy BCG vaccinated or non-vaccinated volunteers not presenting with any clinical symptoms and no previous history of tuberculosis or contact with tuberculosis patients were included in this group.

### Sample Collection

10 ml of peripheral blood was collected from each study subject with prior consent with the help of a trained technician.

### Isolation of peripheral blood mononuclear cells (PBMCs)

Heparinized whole blood was diluted with normal saline in 1:1 ratio and layered on the top of Histopaque (Hi-Media) in 1:1 ratio, followed by centrifugation at 1600 rpm for 30 min. at room temperature. Then upper aqueous plasma layer was discarded, and the buffy coat containing PBMCs was transferred to a new tube. 1 ml of ammonium chloride was added to eliminate any RBCs followed by centrifugation twice at 1600 rpm for 10 minutes, followed by washing with 1X PBS. Isolated PBMCs were used for downstream processing or stored at -80°C until further analysis.

### Isolation of genomic DNA

Genomic DNA (gDNA) from PBMCs was isolated using DNA Isolation Kit (Blood mini kit, Qiagen) as per the manufacturer’s instructions followed by quantification using a nanodrop (Thermofischer scientific).

### Global DNA methylation analysis by COBRA assay

To see if there is any difference in the overall levels of methylation (Global methylation) in TB patients when compared to diseased controls and healthy controls, COBRA assay was performed. 8 samples from each TB and healthy control group and 3 samples from diseased controls were used. 500ng gDNA from each sample was subjected to sodium bisulphite treatment followed by purification of the bisulfite converted DNA using Epitect bisulfite kit (QIAGEN). Bisulfite converted DNA (50ng) was then used for PCR with methylation specific primer of repetitive element i.e. *LINE-1* gene using EpiTaq HS (TakaRa) as per manufacturer’s instructions. Each sample was run as a technical replicate and *LINE-1* gene thus amplified was gel purified using Favorgen Gel purification kit (Qiagen) in accordance to manual provided with the kit followed by restriction digestion using TaqI (Thermofisher scientific) enzyme at 65°C for 1 hour in a PCR thermal cycler (Applied biosystem). Afterwards digested product was run on 2% agarose gel along with 100bp DNA marker followed by densitometry analysis using GelDoc system (Protein simple, alpha view software). For each sample densitometry was repeated three times and average of the same was then taken for final results. The formula used for percent methylation quantification was as follows:

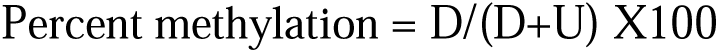

D: Intensity of digested *LINE1* amplicon; U: Intensity of undigested *LINE1*amplicon

### Whole Genome Bisulfite Sequencing and data analysis

Genome-wide DNA methylation patterns of PBMCs from PTB, disease controls and healthy subjects were studied using whole-genome bisulfite sequencing. 4 samples from each group were used. The sequencing libraries were prepared according to the manufacturer’s instructions of ACCEL-NGS® METHYL-SEQ DNA LIBRARY KIT (Swift Biosciences, Inc., Ann Arbor, MI, USA). Briefly, fragmentation of 200ng of genomic DNA was performed using adaptive focused acoustic technology (AFA; Covaris, USA) and the fragmented DNA was bisulfite-treated with EZ DNA Methylation-Gold Kit (Zymo Research, Irvine, CA, USA). These bisulfite-converted DNA fragments were processed for addition of indexing adapters and then purified. The purified libraries were then quantified using qPCR according to the qPCR quantification protocol guide (KAPA library quantification kits for Illumina sequencing platforms) and quality was checked using the Tapestation 4200™ (Agilent technologies). Prepared libraries were sequenced on Novaseq™ (Illumina) at 30X coverage, 2×150bp reads/ sample. Up to 75% of the sequenced bases were of Q30 value >85 meaning very less probability of error highly authentic sequencing data. Sequenced data was processed to generate FASTQ files and proceeded with data analysis.

### Data Analysis for WGB Sequencing

Adapter sequences and poor-quality sequences were removed for paired-end sequence reads with “Trim Galore” tool. Trimmed sequence reads were mapped to reference genome with BS aware alignment tool “bwa-meth”. Next, “MethylDackel” tool was used to detect methylation level at CpG sites. Differential methylation regions were detected with “metilene” and “bsseq” tools respectively. Finally, whole methylome profiling and differential methylation detection was performed with “MethylSeekR” R package. All of the data analysis work was performed by the bioinformatics experts at Wipro Pvt. Ltd. (Bangalore, India).

### Gene Set Enrichment Analysis

For selection of Differentially Methylated Regions (DMRs) a cut off of ± 30% methylation was applied and DMRs qualifying the mentioned criteria were considered for GSE analysis using freely available software GSEA 4.2.2 (Informatics Technology for Cancer Research (ITCR). GSEA-Preranked tool was used for this analysis. First, based on differential methylation analysis, features were ranked and ranking were captured (per cent methylation) in an RNK-formatted file. These Preranked files were used as input in GSEA-Preranked tool and run while choosing Go-Bioprocesses (GOBP) data set from MsigDB (signature database). Positively enriched processes with ≤ 0.25 FDR q-Value and related to immune processes were considered for further analysis. DMRs belonging to the selected processes were then checked for methylation status in promoter region using UCSC genome browser and the ones with differentially methylated promoters were further selected for validation.

### Validation of Whole Genome Bisulfite Sequencing data

Methylation specific primers were designed using Meth Primer 2.0. 50ng of bisulfite converted DNA was used to amplify region of interest by Epitaq HS kit (Takara) in a 50µL reaction. The size of the amplicon resulting from PCR amplification was verified before proceeding further to ensure the correct size.

#### Sanger-sequencing and analysis of DMR of interest

Prepared samples were then sequenced by dye-terminator sequencing. Samples were sequenced on an applied biosystems DNA analyzer with the forward primers of region of interest at a concentration of 5 μM. Sequenced samples from the ABI DNA analyzer were obtained as .ABI files. These files were then analysed using ABI sequence scanner (for PC, available at http://www.appliedbiosystems.com/absite/us/en/home.html). DNA sequences were analysed and peak height was calculated for Cytosine(C) and Thymine(T) in the DNA sequence for calculation of C:T ratio. The sequence files contained the exact same DNA sequence and bp length as the intended DMR amplicon, except for locations at which cytosines were methylated. Cytosine methylation yield two unique chromatogram peaks that corresponded to the presence of cytosine at that position or the presence of thymine at that position. Because bisulfite treatment does not convert methylated cytosines to uracil, the cytosine signal at each CpG location corresponded to the degree of methylation at that site within that sample. Likewise, the thymine signal corresponded to the degree of cytosines that were not methylated (and were therefore converted to uracil during the bisulfite conversion process). Further, the relative size of these peaks was proportionally related to the total percentage of cytosine bases that were methylated. Therefore, methylation levels for each CpG site within the DMR amplicon was quantified by measuring the ratio between peak height values of cytosine (C) and thymine (T), yielding the basic equation for estimation of percent methylation to be (C/(C+T)*100).

### In sillico gene expression analysis of selected genes associated with differentially methylated DMRs in promoter region using Gene Expression Omnibus (GEO2R)

In order to screen the gene expression status of the selected DMR associated genes, 12 GEO transcriptome datasets of PBMCs of TB patients in comparison to healthy and diseased controls were retrieved and reanalyzed using GEO2R tool. GEO2R is a web-based interactive utility that aids users to compare two or more sets of samples from a GEO Series dataset to find differentially expressed genes across varied experimental setups, for example in diseased and healthy condition. In 12 datasets that were retrieved, nine were microarray and rest were RNA sequencing datasets. Using the processed data tables as input, GEO2R performed differential expression analysis using GEOquery and limma (Linear Models for Microarray Analysis) to find genes that exhibit differential expression in a microarray data. For RNA sequencing datasets, a program called DESeq2 (R based) was used to find differentially expressed genes.

---

**Supplementary figure 1:**
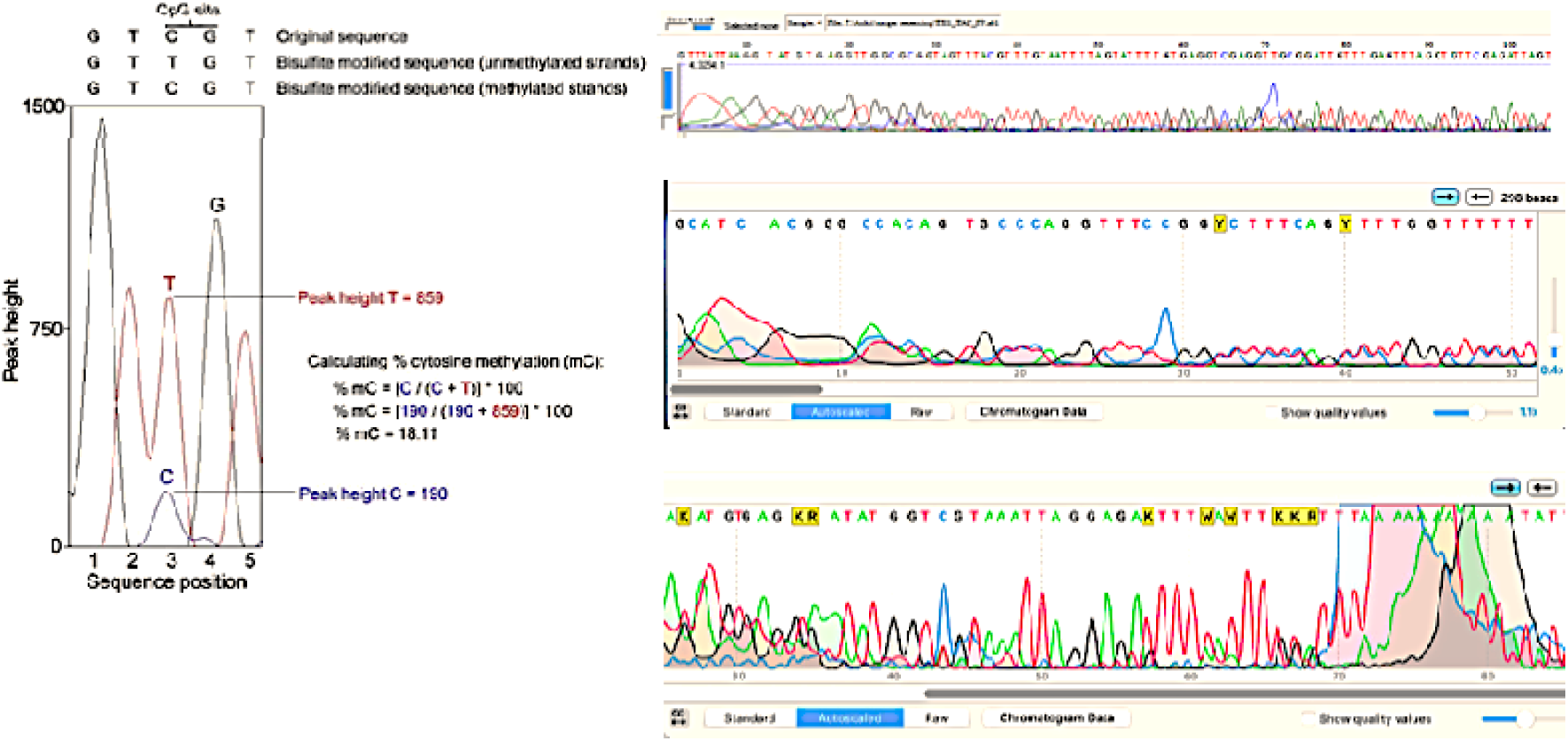
Representative illustration explaining the quantification of percent methylation and representative chromatogram of sanger sequencing

**Supplementary Table 1:**
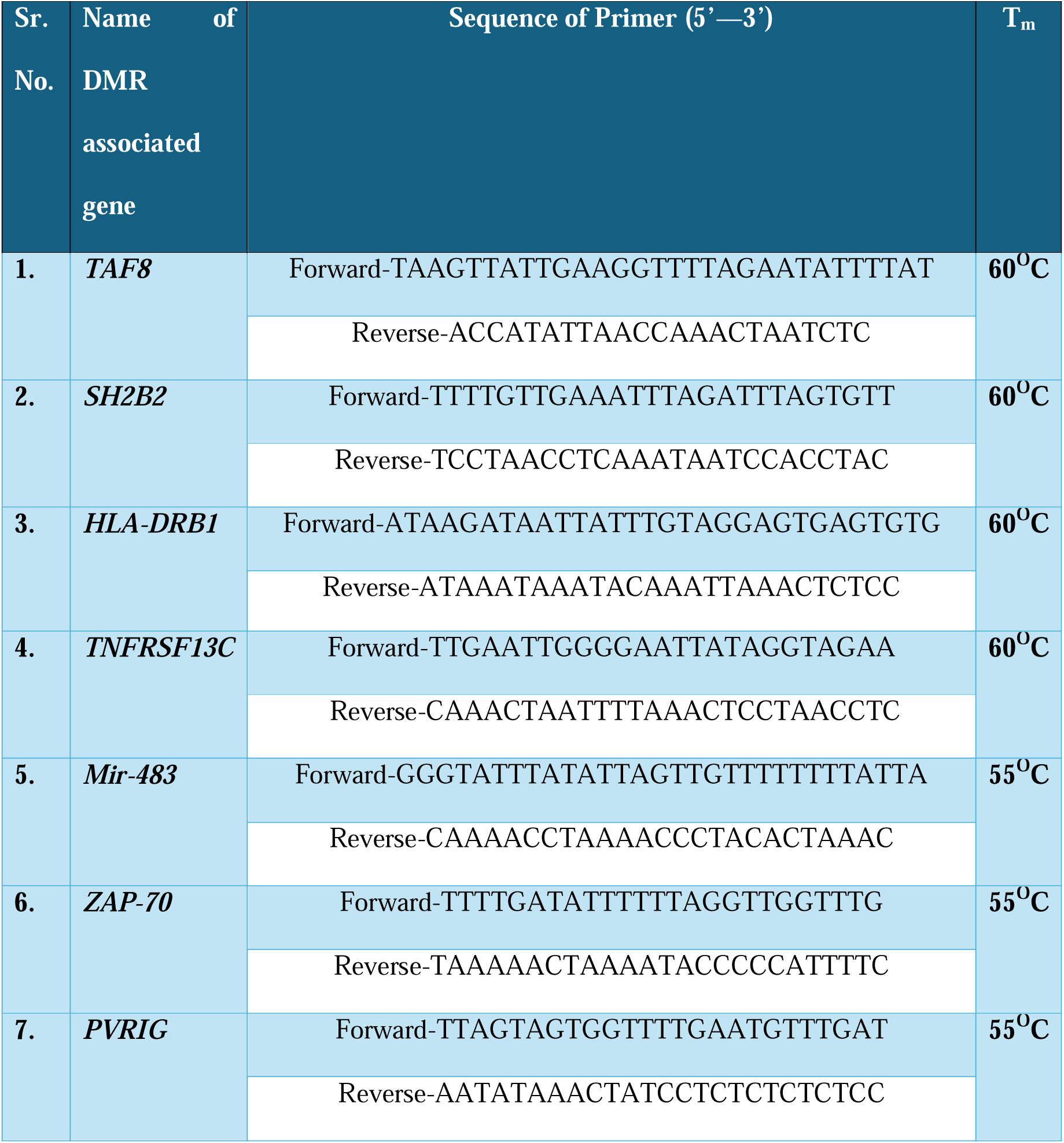
List of methylation specific primers used for DMR validation.

**Supplementary Table 2:** Reaction setup used for Amplification of DMR of interest.

| Reagents | Volume/Amount | Final<br>Concentration/50µl |
| --- | --- | --- |
| TaKaRa EpiTaq HS (5 U/µl) | 0.25 µl | 1.25 U |
| 10X EpiTaq PCR Buffer (Mg<sup>2+</sup> free) | 5 µl | 1X |
| 25 mM MgCl<sub>2</sub> | 5 µl | 2.5 mM |
| dNTP Mixture (2.5 mM each) | 6 µl | 0.3 mM |
| Template | 50ng |  |
| Forward Primer | 20 pmol | 0.4 µM |
| Reverse Primer | 20 pmol | 0.4 µM |
| Nuclease Free water | to 50 µl |  |

**Supplementary Table 3:** Thermal cycling condition used for DMR of interest amplification.

|  | Temperature | Time | Cycles |
| --- | --- | --- | --- |
| **Denaturation** | 98□ | 10 sec | 35 |
| **Annealing** | According to T<sub>m</sub> | 30 sec |  |
| **Extension** | 72□ | 30 sec (upto~ 500 bp) |  |
| **Final extensoion** | 72□ | 10 min | 1 |
| **Hold** | 4□ | ∞ |  |

